# Pre- and postnatal 1,4-phenylene di-isothiocyanate treatment does not induce bile duct injury in neonatal pigs

**DOI:** 10.1101/2025.09.12.675643

**Authors:** Sarah Elefson, Caitlin Vonderohe, Barbara Stoll, Douglas Burrin, Gregory Guthrie

## Abstract

**Background:** Biliary atresia (BA) is the leading cause of pediatric liver transplants, however; the cause of biliary atresia (BA) is unknown. Furthermore, the most common treatment for this disease is with a surgical procedure, which has a greater than 50% failure rate after 5 years. Due to a lack of proper animal models to study the pathology of the disease, little progress has been made in the field.

**Objective:** The objective of this study was to test whether pre and postnatal 1,4-phenylene di-isothiocyanate (DITC) would induce bile duct injury and cholestasis in neonatal pigs.

**Methods:** Pregnant sows received DITC once at gestation week 5 (100 mg/kg; n=2), or 3 times at gestation weeks 5, 6, and 7 (100 mg/kg each; n=2), or twice per week at gestation weeks 5-16 (15 mg/kg each; n=2). Cesarian-delivered piglets of the sows were randomly assigned to receive approximately 200 mg/kg DITC on days two, four, and six of life or to remain untreated. Piglets were fed enterally and collected blood samples were monitored for markers of liver injury for 14 days. At the end of 14 days, tissues were weighed and collected for immunohistochemistry and histopathology scoring.

**Results:** Piglets from sows that received DITC for 11 weeks had lower (*P* < 0.05) final body weight and daily gain compared to other treatments. Piglets from sows that received DITC for 11 weeks had a transient increase in gamma-glutamyl transferase. Liver histological scoring and analysis also did not show signs of BA. Piglets that received DITC from 11-week DITC-treated sows had elevated hepatic bile acids (*P* < 0.05), but there was no difference in serum bile acids (*P* > 0.05).

**Conclusions:** The administration of DITC to pregnant sows and neonatal piglets did not result in the development of bile duct or hepatic injury.

## 1. Introduction

Biliary atresia (BA) occurs in 1 of every 12,000 live births (2) in the United States, and globally, affects between 0.5 and 1.6 in 10,000 live births (3, 4). The signature characteristic of BA is atresia or the absence of the common bile duct. When BA occurs, both extra- and intra-hepatic bile ducts are impacted, resulting in obstruction of bile flow. BA manifests with hepatic inflammation and fibrosis, but the etiology of BA is unknown (5) Consistent among all forms of BA is the increase in serum direct/conjugated bilirubin, due to the near-total extrahepatic obstruction of bile flow (6). Additionally, typical clinical pathology markers of cholestasis and injury, including serum gamma glutamyl transferase (GGT), alanine amino transferase (ALT), and aspartate amino transferase (AST) are elevated (7). Given then clinical markers of BA are usually detected in early postnatal life (8), it has been postulated that BA pathogenesis originates during pregnancy and manifests after birth.

The most common treatment for this disease is a surgical procedure, called hepatoportoenterostomy (HPE), where the proximal duodenum is surgically attached to the liver, providing a means for bile to be delivered directly to the intestines, bypassing the damaged extrahepatic bile ducts (9). Unfortunately, the success rate of HPE is about 50% after 2 years, so many children will require a liver transplant (10). Developing new clinical strategies to treat symptoms of BA and improve outcomes pre- and post-HPE is difficult due to the low incidence of the disease. There has been limited progress in developing animal models to phenocopy damage to the extrahepatic biliary system. In mice, models of seen with BA murine models of BA include most frequently have used rhesus rotavirus infections (11) and bile duct ligation (12), with less frequent approaches such as global knockout of Sox17 (13). Additionally, toxin models have been developed including the use of bilatresone in zebrafish (14), and 1,4-phenylene di-isothiocyanate (DITC) in rats (15) and pigs (16) to differing successes.

Establishing a consistent and pathologically comparative animal model to study this disease is essential to be able to investigate post-HPE adaptation and identify novel targets for intervention. Thus, a neonatal animal model that replicates the bile duct injury found in BA, and is large enough for HPE, would permit the studying of the physiological and molecular adaptations that occur post-HPE. Piglets are an excellent model for studying human diseases as they are comparable in size to human infants, and have similar anatomy and physiology (17, 18). The piglet has previously been used to study bile duct injury and disease (19-21), and one study had reported the use of DITC (16) to induce BA in piglets during sow gestation; however, the findings from these studies were equivocal. Therefore, the objective herein is to establish a piglet model using DITC treatment during gestation, imitating the pathology of developmental BA in the piglets.

## 2. Materials and Methods

All experiments were approved by the Animal Care and Use Committee of Baylor College of Medicine. 1,4-Phenylene di-isothiocyanate (DITC) was purchased from TCI (Tokyo Chemical Industries; Tokyo, Japan). Pregnant sows (N=6) received DITC either once (Sow1; n=2; 100 mg/kg DITC on gestational week 5), three times (Sow3; n=2; 100 mg/kg DITC on gestational weeks 5, 6, and 7), or 11 weeks (Sow11; n=2; 15 mg/kg DITC starting on gestational week 5, twice per week, through the end of gestation). Piglets were delivered by cesarian section from sows on gestation d 113 (out of 114) and surgically implanted with a jugular catheter and an orogastric tube. Piglets were randomly assigned to be untreated (Sow#-CON) or received approximately 200 mg/kg DITC on postnatal d 2, 4, and 6 (Sow#-DITC). Sows received DITC mixed with 200 g of sugar and piglets received DITC mixed with pig milk replacer (Litterlife; Merrick, Middleton, WI).

### 2.1 Serum Chemistry

All blood samples were centrifuged at 2,000 g for 10 minutes at 4°C to separate the serum from the red blood cells. Serum was stored at -80°C until time of analysis. Serum chemistry samples were analyzed by the Comparative Pathology Laboratory at Baylor College of Medicine using commercially available kits with a Cobas Integra 400 plus analyzer (Roche Diagnostics, Rotkreuz, Switzerland) (22).

### 2.2 Tissue Collection

Animals were humanely euthanized on d 14 using a commercially available pentobarbital solution (Euthasol; Virbac Animal Health, Carros-Cedex, France). Tissues were weighed before sub-sampling. The liver flash frozen in liquid nitrogen and stored at -80°C until time of analysis and fixed in 10% neutral buffered formalin. Tissues were then embedded in paraffin and slides were cut. Staining for H&E, Sirius Red, and Pan Cytokeratin (DAKO, Agilent, M3515) was performed at the Texas Medical Center Digestive Disease Center Pathology Core.

### 2.3 Bile Acid Quantification

Bile acid concentrations in the plasma and liver were analyzed as previously described in Vonderohe et al. (23), using a commercially available kit (Genway, San Diego, CA).

### 2.4 Liver Histopathology Quantification

Liver sections were stained with Sirius Red to measure hepatic fibrosis. Fiji Image J (24) was utilized to quantify the intensity of Siris Red staining on each slide. Briefly, color channels on the histology slide image were split to isolate the red color channel, then mean amount of red was measured for the entire histology slide to obtain the mean amount of red that was present.

Liver sections were immunohistochemically stained with pan cytokeratin (DAKO, Agilent, M3515, includes cytokeratin 7 and cytokeratin 19). QuPath 4.3 (25) cell detection was used to measure the number of cells that were stained positively with 3,3′-Diaminobenzidine (DAB). To quantify the number of cells that were DAB positive, the following criteria were set: intensity parameter threshold of 0.1, max background intensity of 2, and intensity threshold parameters with a score compartment of Nucleus: DAB optical density mean, threshold 1+ = 0.25, threshold 2+ = 0.3, and threshold 3+ = 0.4.

### 2.5 Liver Histology Scoring

Liver slides stained with hematoxylin and eosin were submitted to Texas A&M University’s Veterinary Medicine and Biomedical Sciences department. An American College of Veterinary Pathologist-boarded veterinary pathologist, blinded to the treatments, scored the slides. The slides compared to tissue from studies where piglets were enterally fed for 2 weeks without receiving DITC to tissue collected at the end of the study. Liver sections were scored for inflammation, ductular reaction, bile stasis, and liver pallor. All categories were scored on a scale of 0-3 where 0 indicated the absence of the parameter in question and 3 represented a severe presence of the parameter.

## 3. Statistical Analysis

Data were analyzed in SAS 9.4 and graphed in GraphPad Prism 10.1.1. All growth data, serum chemistry, histology measurements and scores were analyzed using a nested and mixed model in which piglet treatment was nested within sow treatment. Fixed effects included treatment, sex, and time, and a random effect of piglet. Repeated measures were utilized for serum parameters. If a parameter had a treatment by time interaction, multiple comparisons within specific days were analyzed individually to further identify when differences occurred throughout the study. *P*-values < 0.05 were considered significant and *P*-values ≤ 0.10 were considered a trend.

## 4. Results

### 4.1 Growth and Relative Organ Weights

We were the first to measure the impact of DITC administration on piglet growth, as previously solely liver histopathological parameters were reported. There was a trend (*P* = 0.05) in birth weight across treatments (**Figure 1.A**), where piglets born from Sow3-DITC had a heavier average body weight than those from Sow1-CON (1,718 g vs 1,160 g, respectively). At the end of the study, there was a sex difference (*P* = 0.04) where males weighed more than females (2,739.74 g vs 2,307.34 g, respectively). Additionally, piglets born from Sow3-CON and Sow3-DITC weighed more (*P* < 0.01) than Sow11-CON and Sow11-DITC (3,216.67 g and 3,328.13 g vs 1,853.87 g and 1,653.25 g, respectively; **Figure 1.B**). Consequently, piglets from Sow11-CON and Sow11-DITC gained less (*P* < 0.01) weight than the other treatment groups (**Figure 1.C**). Relative weights for the liver and heart were highest (*P* < 0.01) for the Sow11-DITC piglets compared to other treatments (**Figure 1.D and Figure 1.E**). Also, females had a heavier relative heart weight (*P* < 0.01) compared to males (8.5 g/kg bodyweight vs 6.8 g/kg body weight, respectively; sex effect data not shown). Distal ileum and overall small intestinal relative weights were lower for the Sow11-DITC piglets compared to the Sow3-CTL (19.21 g/kg BW vs 26.39 g/kg BW and 36.43 g/kg BW vs 44.20 g/kg BW, respectively; **Figure 1.G and Figure 1.H**).

**Figure 1.**
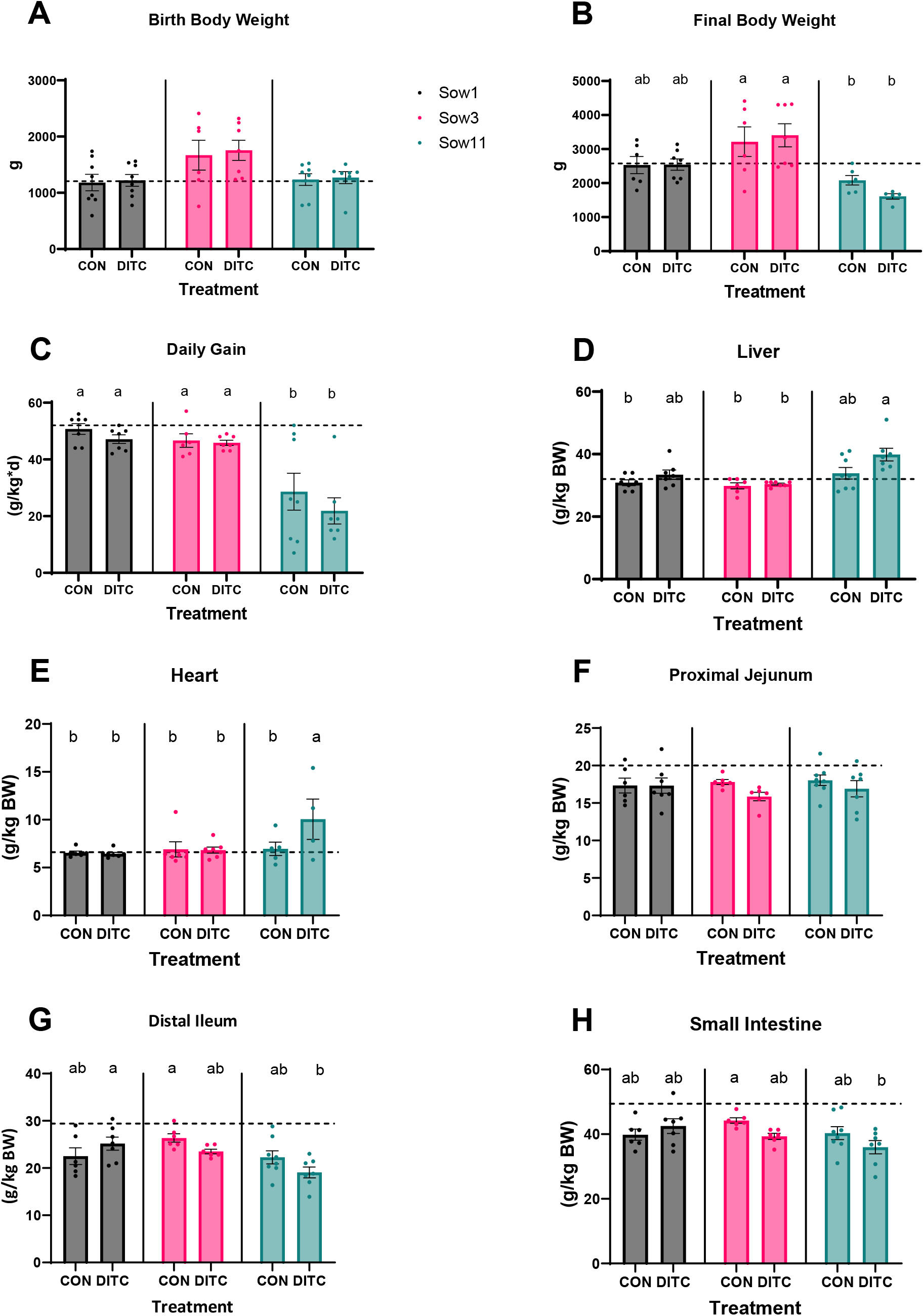
Body and relative organ weights. A) Birth body weight of pigs at the start of the study. B) Final body weight of pigs at the end of the study. C) Average daily gain of pigs throughout the study. D-H) Organ weights relative to body weight on d 15. Differing superscripts indicate statistical differences of P < 0.05. CON, piglet control; DITC, piglet received 1,4-phenylene di-isothiocyanate; Sow#, # is the number of times sows received 1,4-phenylene di-isothiocyanate. Dotted line represents historical controls at d 15.

### 4.2 Serum Chemistry

We also examined whether DITC treatment affected standard serum markers of cholestasis and liver injury. Overall, there was a time effect (*P* < 0.05) for all serum parameters (**Figure 2**) where serum blood urea nitrogen (BUN; **Figure 2.A**), alkaline phosphatase (ALP; **Figure 2.B**), and gamma glutamyl transferase (GGT; **Figure 2.C)** increased between birth and d 14. The BUN levels were elevated for Sow11-CON and Sow11-DITC compared to all other treatment groups. ALP values were highest for Sow1-CON piglets compared to all other treatment groups. Gamma glutamyl transferase values were the highest for Sow11-DITC and Sow1-DITC compared to all others.

**Figure 2.**
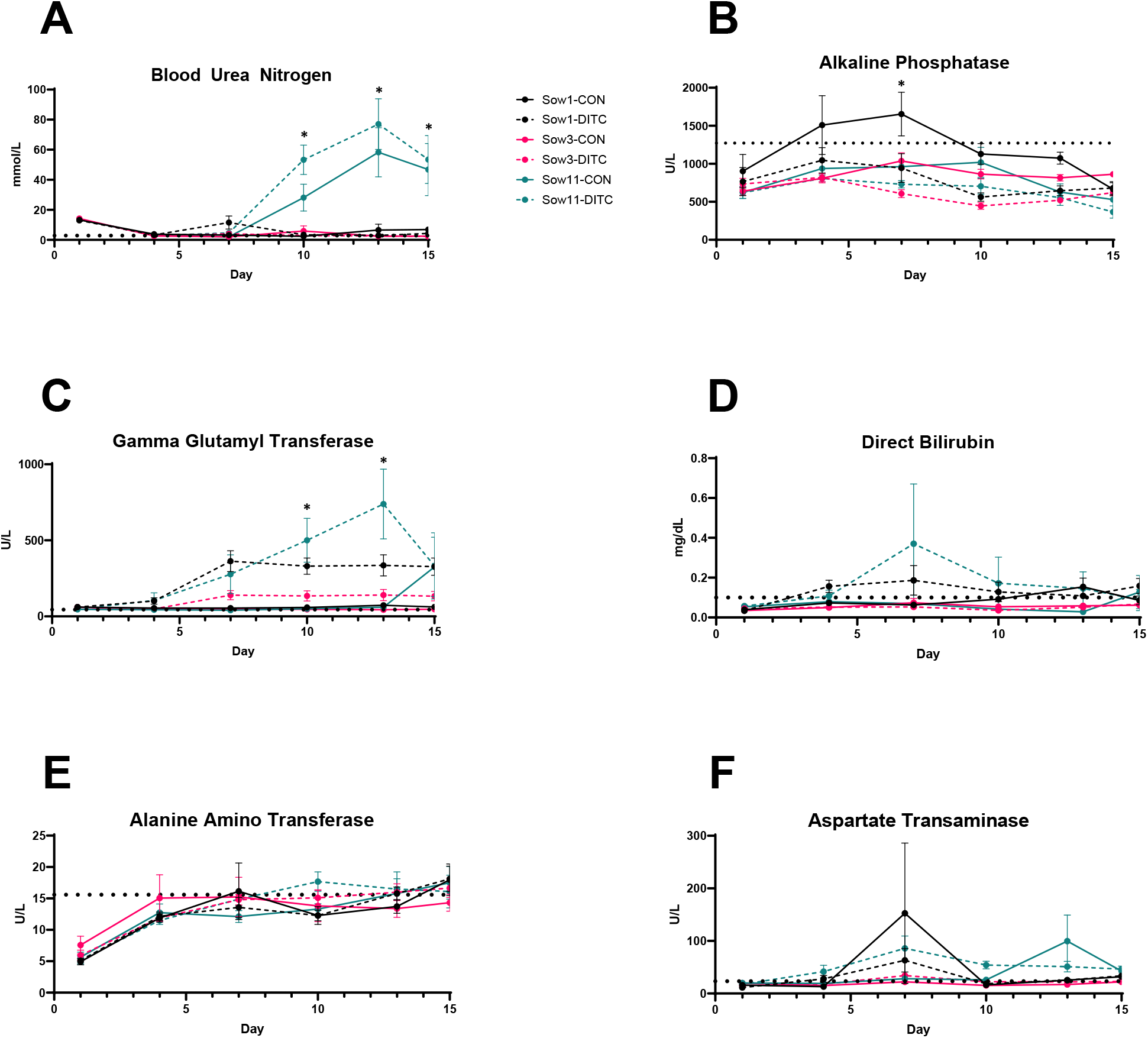
Serum chemistry of piglets starting d 1 of the study through d 15. A * located above a treatment group indicates statistical difference (P < 0.05). CON, piglet control; DITC, piglet received 1,4-phenylene di-isothiocyanate; Sow#, # is the number of times sows received 1,4-phenylene di-isothiocyanate. Dotted line represents historical controls at d 15.

There was a statistically significant interaction between treatment and time (*P* < 0.05) for BUN and GGT, and a tendency (*P* < 0.10) for ALP. These interactions are indicative of transient effects of the treatments over the course of the study. On d 10 of the study, BUN levels were significantly elevated and stayed elevated for Sow11-CON and Sow11-DITC piglets compared to other treatment groups throughout the remainder of the study. On d 10 and d 13 Sow11-DITC pigs had elevated GGT levels compared to all treatments except Sow1-DITC. However, this did not persist through d 15 of the study. ALP was elevated for Sow1-CON on d 7 compared to all treatments except Sow3-CON. Sow1-CON was also the most elevated on d 10 but was only statistically different from Sow3-DITC. This difference did not persist, on d 15 Sow3-CON was the most elevated, and statistically different from Sow11-DITC. There were no treatment by time effects for direct bilirubin (**Figure 2.D**), alanine amino transferase (**Figure 2.E**), and aspartate transaminase (**Figure 2.F**).

### 4.3 Bile Acids

Plasma total bile acid concentrations were similar across all treatment groups on d 14 (*P* > 0.05; **Figure 3.A**). However, hepatic total bile acids were statistically different (*P* = 0.01) on d 14. Total hepatic bile acids concentrations (**Figure 3.B**) were lowest in Sow1-DITC (86.00 µmol/g) and increased in Sow11-DITC and Sow3-CON (212.32 µmol/g and 203.12 µmol/g, respectively) (*P* = 0.01).

**Figure 3.**
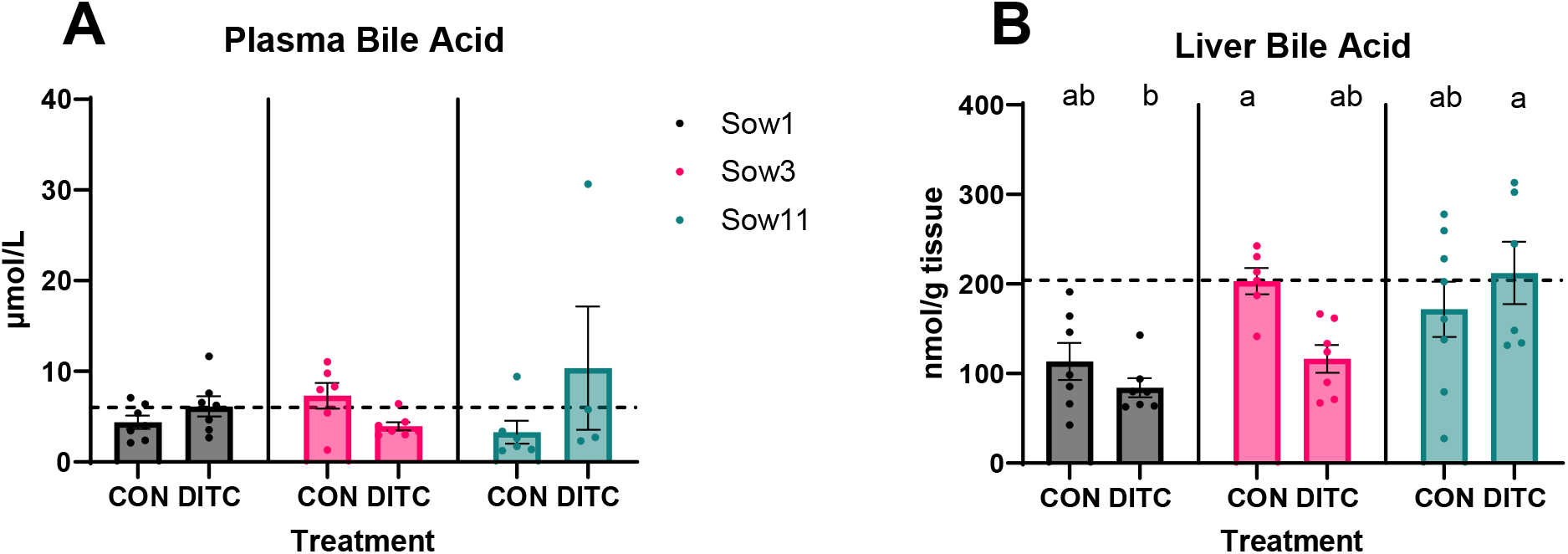
Bile acids. A) Plasma and B) liver bile acids at d 15 of the study. Differing superscripts indicate statistical differences of P < 0.05. CON, piglet control; DITC, piglet received 1,4-phenylene di-isothiocyanate; Sow#, # is the number of times sows received 1,4-phenylene di-isothiocyanate. Dotted line represents historical controls at d 15.

### 4.4 Hepatic Injury

In addition to serum markers of cholestasis, we also examined what impact DITC treatment had on hepatic injury markers of fibrosis and ductular reaction. No differences (*P* > 0.05) were found between the quantitative measures of Sirius red intensity (**Figure 4.A**). Representative images of Sirus Red staining are depicted in **Figure 4.B**. There were also no differences (*P* > 0.05) in the percentage of cells positively stained for pan cytokeratin across treatments (**Figure 4.C)** Representative images of pan cytokeratin staining are depicted in **Figure 4.D**.

**Figure 4.**
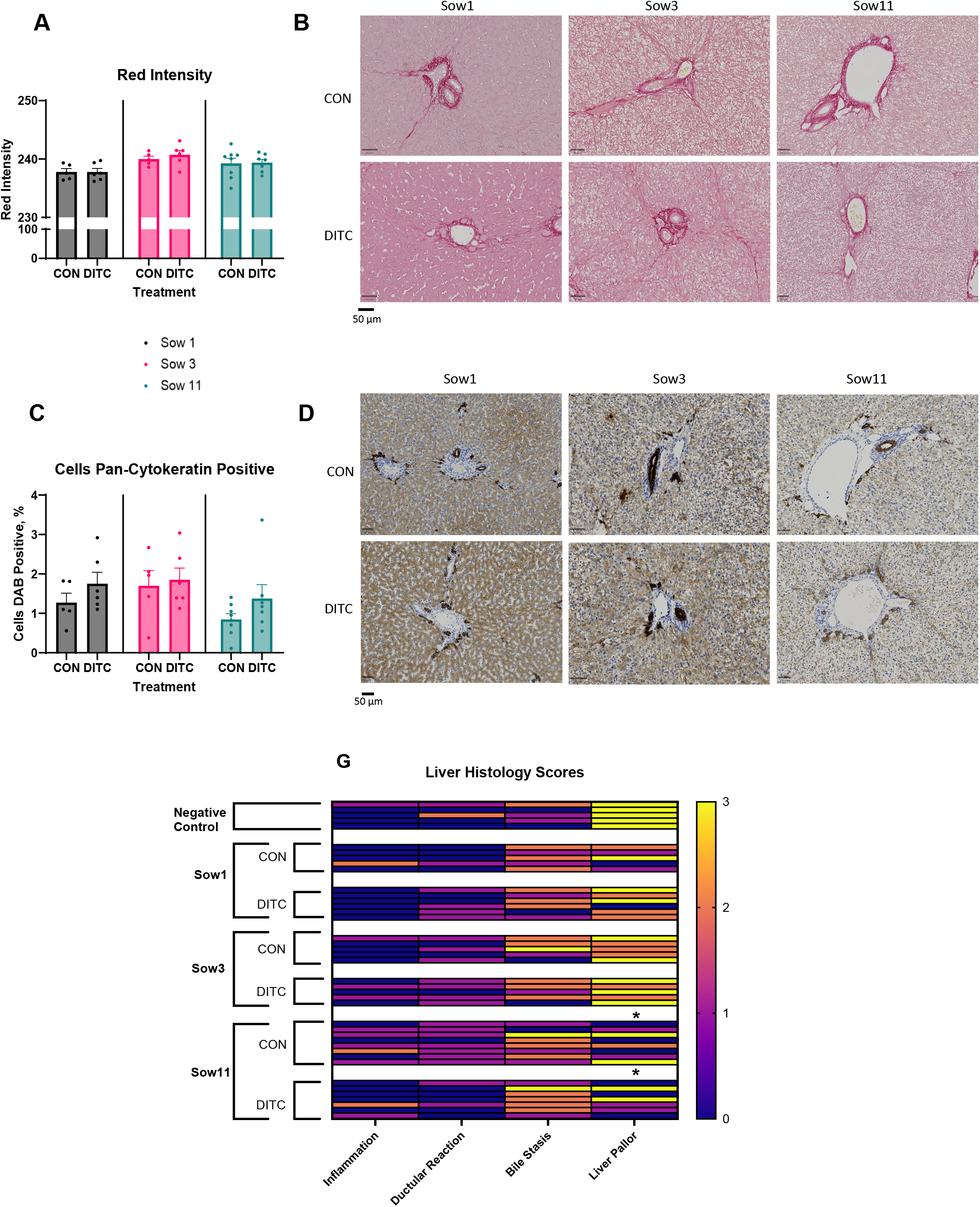
Liver staining and histology. A) Average amount of red from Sirus Red staining measuring fibrosis. B) Representative image of Sirus Red staining from each treatment. E) Percent of cells identified through 3, 3’-diaminobenzidine staining with a pan-cytokeratin antibody. C) Representative image of pan-cytokeratin staining from each treatment. E) Histological scoring for liver disease. CON, piglet control; DITC, piglet received 1,4-phenylene di-isothiocyanate; Sow#, # is the number of times sows received 1,4-phenylene di-isothiocyanate.

Based on inflammation, ductular reaction, bile stasis, and liver pallor, there was a difference (*P* = 0.02) in liver pallor on slides collected from Sow11-CON and Sow11-DITC when compared to control slides collected from pigs that never received DITC (1 and 1 vs 3, respectively, **Figure 4.G)**. This indicates that piglets that never received DITC had more pale livers than piglets that received DITC. The clinical relevance of this is unclear. No other differences (*P*> 0.05) were found across the scored categories.

### 5.1 Discussion

Animal models to accurately and consistently replicate the clinical presentation and pathogenesis of biliary atresia are needed to study interventions and treatments for infants with biliary atresia. Previous animal studies have suggested the use of the compound, DITC, can cause an atresia-like phenotype in rats and pigs, though the number of studies is very sparse and supporting data limited. Thus, in this study, we evaluated whether treatment with DITC in both pregnant sows and neonatal piglets would result in newborn piglets exhibiting an atresia-like phenotype, thereby assessing the drug’s efficacy.

The mechanism by which DITC may cause ductular injury has not been elucidated; however, similar compounds that contain isothiocyanate, like α-naphthylisothiocyanate (ANIT), have also been shown to cause damage to extrahepatic bile ducts via oxidative stress and inflammation (26-28). ANIT induces free radical oxidative stress resulting in conjugation and depletion of the glutathione (GSH) pool leading to biliary epithelial injury (29). In the cholangiocyte, pH-dependent deconjugation occurs leading to elevated free ANIT, which may recruit neutrophils (30). The resultant effect is detection of high ROS and inflammation, primarily in the cholangiocytes (31, 32). In our study, we observed a transient significant increase in GGT concentrations following treatment with DITC in our Sow11-DITC group and a trend for increased GGT in the Sow1-DITC group. GGT is a marker of cholangiocyte injury, this suggests that DITC is causing injury through a similar mechanism as ANIT. However, this injury was not sustained and all groups had similar GGT by the completion of the study. In the context of BA, this biliary epithelial injury was not consequential to bile flow since there was no change in serum direct bilirubin levels. The piglets may have also experienced some off-target injury from DITC toxicity, as well, given that blood urea nitrogen was elevated, which may indicate kidney damage or reduction in functional mass. However, it is important to note that all serum levels were within appropriate ranges for pigs as listed by the Iowa State University, College of Veterinary Medicine Clinical Pathology Reference Intervals (33).

Another characteristic of cholestasis in pigs with other forms of obstructive cholestasis is the elevation of serum and hepatic bile acids (34). Despite Sow11-DITC and Sow3-CON having elevated hepatic bile acid concentrations, there were no differences in the serum bile acid concentration across treatments. Furthermore, the serum bile acid levels we reported are comparable to enterally fed pigs without liver injury (35-37). Previously, it was observed, mainly through liver histology scoring, that progeny from sows who received DITC during the 5^th^ week of gestation, showed clinical signs of BA (16). However, Lainakis et al. (16) also reported that there were no other clinical signs of BA being present in their pigs administered DITC. Our histopathology findings do not replicate the results described in Lainakis et al. There was a significant lack of histological markers that would be indicative of injury to the bile ducts, either through atresia of the intrahepatic ducts, or of ballooning of extrahepatic ducts from distal blockage of bile flow.

Additionally, other histology markers of hepatic injury were not different among groups. Sirus Red was used to measure fibrosis in the liver; it is important to note that pig livers are naturally fibrous (38). Thus, this parameter relied on differences that could be attributed to treatments. However, there were no differences in the amount of fibrosis measured through Sirus Red staining across treatments. An increase in ductular reaction, characterized by the proliferation of cholangiocytes typically around the portal vein, is commonly observed during biliary atresia (BA) and other forms of obstructive cholestasis (39). We used pan cytokeratin staining to measure bile duct development and quantify the degree of ductular reaction in these piglets, but there were no clinically or statistically significant differences across treatments.

Despite a lack of injury to the livers of piglets, we believe there was a systemic effect of pre-natal DITC exposure. Piglets that were exposed to DITC for 11 weeks in utero had a lower average daily body weight gain compared to piglets that had fewer exposure times. This occurred in both piglets that were fed control or DITC-supplemented diets, suggesting the effect is from in-utero exposure and not post-delivery exposure to DITC. Our piglets experienced gastrointestinal distress, including diarrhea and vomiting when exposed to post-natal DITC in the diet. This finding is supported by reports dating from the early 1970s that DITC has side effects in humans including diarrhea, nausea and vomiting (40), with the incidence in some cases ranging from 30-50% (41, 42). Thus, it was not unexpected that the piglets had similar side effects, particularly vomiting and diarrhea, which may be from direct effects on the GI tract. However, based on our findings, this does not seem likely to be due to liver disease.

## 6. Conclusions

In summary, in contrast to a previous report (16), our results show that treatment with DITC in pregnant sows and neonatal piglets does not cause profound hepatic injury or cholestasis. Our findings are based on DITC treatment using the previously reported dose, as well as conditions involving increased dose exposure. Furthermore, varying amounts of DITC were utilized to observe a potential dose-response for the development of fibrotic cholangitis, but this was not the case. We also showed that DITC treatment caused mild clinical signs of toxicity in neonatal pigs such as diarrhea, uremia, and reduced growth rate, it is not an effective chemical approach to a viable or reproducible study BA.

## Author Contributions: Sarah Elefson

Formal analysis, Investigation, Data curation, Writing – original draft; **Caitlin Vonderohe**: Data curation, Investigation, Editing; **Barbara Stoll:** Data curation; Investigation, Methodology, Investigation, Editing; **Douglas Burrin:** Investigation; Funding acquisition, Methodology, Project administration; Supervision, Resources, Editing; **Gregory Guthrie:** Conceptualization, Data curation, Funding acquisition, Investigation, Methodology, Project administration, Supervision, Writing – reviewing and editing

## Acknowledgments

Special thanks Liwei Cui, Inka Didelija, and Xiaoyan Chang for their assistance in sample analysis. Appreciation is also extended to Dr. Yava Jones-Hall at Texas A&M for liver histological scoring.

